# Molecular dynamic simulations to investigate the structural impact of known drug resistance mutations on HIV-1C Integrase-Dolutegravir binding

**DOI:** 10.1101/781120

**Authors:** Rumbidzai Chitongo, Adetayo Emmanuel Obasa, Sello Given Mikasi, Graeme Brendon Jacobs, Ruben Cloete

**Affiliations:** South African Medical Research Council Bioinformatics Unit, South African National Bioinformatics Institute, University of the Western Cape, Cape Town, South Africa; Division of Medical Virology, Department of Pathology, Faculty of Medicine and Health Sciences, Stellenbosch University, Tygerberg, Cape Town, South Africa

## Abstract

Resistance associated mutations (RAMs) threaten the long-term success of combination antiretroviral therapy (cART) outcomes for HIV-1 treatment. HIV-1 Integrase (IN) strand transfer inhibitors (INSTIs) have proven to be a viable option for highly specific HIV-1 therapy. The INSTI, Dolutegravir is recommended by the World Health Organization for use as first-line cART. This study aims to understand how RAMs affect the stability of IN, as well as the binding of the drug Dolutegravir to the catalytic pocket of the protein. Molecular modelling of HIV-1C IN was performed using the SWISS-MODEL webserver; with quality assessment performed using internal methods and external software tools. The site directed mutator webserver was used to predict destabilizing and/or stabilizing effects of known RAMs while FoldX confirmed any changes in protein energy upon introduction of mutation. Interaction analysis between neighbouring residues was done using PyMOL. Three randomly selected mutations were chosen for molecular dynamic simulation studies using Gromacs. Trajectory analysis included Root mean square deviation and fluctuation, Radius of gyration, Principal component analysis and Interaction analysis between Dolutegravir and protein residues. The structural quality assessment indicated high reliability of the HIV-1C IN tetrameric structure, with more than 90% confidence in modelled regions. Change in free energy for the G140S mutant indicated a stabilizing effect and simulation analysis showed it to affect structural stability and flexibility of the protein structure. This was further supported by the drug being expelled from the G140S mutant active site, as indicated by interaction analysis. Our findings suggest the G140S mutant has a strong effect on the HIV-1C IN protein structure and Dolutegravir binding and should be validated using laboratory-based experiments. This approach can be applied to determine the effect of other mutations/variants on HIV-1C integrase drug binding.

## Introduction

The Integrase (IN) enzyme plays an important role in the Human Immunodeficiency Virus type 1 (HIV-1) replication cycle by catalysing two distinct reactions termed: 3’-end processing and strand transfer. During the 3’ processing, IN removes two nucleotides from the 3’ ends of both viral DNA strands and exposes the C-alpha hydroxyl group on the 3’ends. The subsequent step involves strand transfer whereby, IN attacks the phosphodiester backbone of the host DNA and links the exposed 3’-end to the 5’ hydroxyl end of the host DNA [1]. This makes HIV-1 IN an important target for combination antiretroviral therapy (cART). HIV-1 IN is a 32 kilo Dalton (kDa) protein, and consist of three structural and functional domains; the N-terminal domain (NTD, residues 1-49), the catalytic core domain (CCD, residues 50-212), and C-terminal domain (CTD, residues 213-288). It also contains a conserved DDE motif consisting of residues Asp54, Asp116 and Glu152 in the CCD, important for drug binding and enzyme activity [2]. Several IN strand transfer inhibitors (INSTIs) have been developed [3–5]. These inhibitors include; Raltegravir (RAL) and Elvitegravir (EVG) as first-line INSTIs and the most recently approved second-line inhibitors Dolutegravir (DTG) and Bictegravir (BIC) [6]. DTG is a coplanar and linear molecule with a high barrier against resistance, it is safe and tolerable and shows little to none drug interactions [7]. Furthermore, a recent study provided evidence for the replacement of RAL with DTG based on the low prevalence of DTG resistance and the low risk for INSTI mutations when patients are on DTG treatment [8]. Although BIC has been recently approved, not much information is known about it and as a result, DTG still remains the preferred option.

The functional mechanism of INSTIs is to bind to the catalytically essential magnesium ions, thereby displacing the reactive 3’-hydroxyl group of the terminal A17 away from the active site which disrupts the strand transfer process. Several mutations have emerged in patients receiving first-line INSTIs, RAL and EVG. Brado *et al*. reported that despite higher fold RAMs against INSTIs being absent in most treatment naïve patients, they can emerge under treatment, particularly with first generation INSTIs [9]. Several studies have used the prototype foamy virus intasome structure (medium sequence identity) to model the structure of HIV-1 IN in order to investigate the effect of mutations on HIV-1 IN using molecular dynamic simulations [10]. These studies revealed the binding mode of EVG and RAL to HIV-1 IN and the structural mechanism of drug resistant mutants that affect the 140’s loop region spanning residues 140-149. The precise role of this loop is unknown, but molecular dynamics studies have demonstrated the importance of this loop’s flexibility for catalysis [11]. This loop has been reported to regulate the active site of HIV-1 IN by decreasing or increasing flexibility under the influence of mutations G140A and G149A. Conformational flexibility of this loop is thought to be important for the catalytic steps following DNA binding, as a decrease in flexibility induced, for example, by the G140A mutation, results in lower levels of activity, despite minimal effects on DNA binding [11]. Mouscadet *et al*. [12] reported that the G140 residue is not directly involved in the cooperative flexibility of the catalytic loop, but plays a critical role in controlling the overall motion of the loop and its precise position relative to the phosphodiester bond to be cleaved. The G140 residue also participates in the catalytic loop hinge formation and its mutation may restore specific contacts required for catalysis, between the loop of the double mutant and the end of the viral DNA [11,13,14]. The findings from these studies were inconclusive due to the poor quality of the protein models delineating the active site and viral DNA binding site for simulation studies.

In 2017, Cryogenic electron microscopy was used to solve the structure of HIV-1 strand transfer complex intasome for HIV-1 subtype B [15]. This provided us with a unique opportunity to model the structure of HIV-1 subtype C IN to interrogate the effect of known drug resistance associated mutations (RAMs) on the protein structure using molecular dynamic simulation studies. This is the first study that uses the consensus wild type subtype C IN sequence to build an accurate 3D model of HIV-1C IN to understand the effect of three known mutations on DTG drug binding.

## Materials and Methods

### Generation of consensus HIV-1C Integrase sequence

To compare our sequences with the rest of the IN sequences from South Africa, we performed a search on the HIV Los Alamos National Library (LANL) database (https://www.hiv.lanl.gov/components/sequence/HIVsearch.com). Our search inclusion criteria included all South African HIV-1 subtype C IN sequences and those identified from treatment naïve patients. We selected one sequence per patient and all problematic sequences were excluded from further analyses. Finally, the consensus sequence was generated using the database-derived HIV-1C_ZA_ sequences (**n = 314**) and cohort sequences (**n = 91**) [9]. Nucleotide sequences were verified for stop codons, insertion and deletions using an online quality control program on the HIVLANL database (https://www.hiv.lanl.gov/content/sequence/QC/index.htm). Multiple sequence alignments were done with MAFFT version 7, from which the consensus sequence was derived [16]. As part of quality control, each of the viral sequences were inferred on a phylogenetic tree in order to eliminate possible contamination.

### Molecular modelling and quality assessment

The crystal structure of the HIV-1B intasome (PDBID: 5U1C) was used to generate a three-dimensional tetrameric structure of HIV-1C IN using the consensus HIV-1C sequence that we generated. The SWISSMODEL webserver was used for model generation by first constructing a pairwise sequence-structure alignment between HIV-1C wild-type (WT) amino acid sequence and template 5U1C [17]. The quality of the resulting model was assessed using SWISSMODEL quality assessment scores, Root mean square deviation analysis compared to homologous template (PDBID: 5U1C) and with publicly available algorithms located at the SAVES webserver (https://servicesn.mbi.ucla.edu/SAVES/) namely; ERRAT, VERIFY3D and Ramachandran plot [18,19].

### Structure preparation

The predicted 3D structure of HIV-1C IN was superimposed to 5U1C to extract proviral DNA, while the Magnesium (MG) ions and drug DTG were obtained from homologous template 3s3m (Prototype foamy virus). The wild-type (WT) structure of HIV-1C was energy minimized in complex with DNA, MG and DTG using Gromacs version 5.1 [20]. Thereafter, we predicted the stabilizing and/or destabilizing effect of mutations on the protein structure. For this purpose, the site directed mutator (SDM) webserver and the software FoldX was used to predict the change in Gibbs free energy after the introduction of the mutation. We also calculated the loss or gain of polar interactions between neighbouring residues located adjacent to the mutation using PyMOL [21].

### Molecular dynamic simulation

For simulation studies we only considered the two inner dimers of the protein structure, as the other two monomers were similar in sequence and structure. Three different mutant systems were prepared by introducing a specific mutation into the WT structure through the mutagenesis wizard in PyMOL and energy minimizing the structures using Gromacs [20]. The WT and three mutant systems (E92Q, S140 and R143) were prepared by uploading the atomic coordinates of the Protein-DNA-MG-DTG complexes to CHARMM-GUI interface [22]. The three mutant systems selected for simulation studies represent three resistance pathways associated with RAL, EVG and possibly DTG resistance. The option solution builder was used as an input generator. Each system was solvated in a rectangular TIP3 water-box with 10Å distance between the edges of the box. The topology and coordinates for each system was generated using CHARMM36 force field and CHARMM general force field for DTG. Each system was neutralized by adding counter ions to each of the systems. For the WT system, 157 potassium ions (K) and 81 chloride ions (Cl) were added, for the mutant R143 system we added 156 K and 81 Cl ions, while for the mutant system S140 we added 157 K and 81 Cl ions and for the mutant system Q92 we added 335 K and 276 Cl ions. Each system was at a final concentration of 0.15M for simulation dynamics.

Gromacs version 5.1 was used for running all the simulations [20]. Each system underwent 50000 steps of steepest descents energy minimization to remove steric overlap. Afterwards, all the systems were subjected to a short position restraint NPT (constant Number of particles, Pressure and Temperature) for 500 picoseconds (ps) to stabilize the pressure of the system by relaxing the system and keeping the protein restrained. For NPT, the Nose-Hoover pressure coupling was turned on with constant coupling of 1ps at 303.15K under conditions of position restraints (h-bonds) selecting a random seed. Electrostatic forces were calculated using Particle Mesh Ewald method [20]. All systems were subjected to a full 300 ns simulation under conditions of no restraints.

The analyses of the trajectory files were done using GROMACS utilities. The root mean square deviation (RMSD) was calculated using gmx rmsd and root mean square fluctuation (RMSF) analysis using gmx rms. The radius of gyration was calculated using gmx gyrate to determine if the system reached convergence over the 300 nanoseconds (ns) simulation. Pairwise distance analysis between the drug and MG was done using gmx pairdist tool. Afterwards we extracted structures every 50 ns over the last 200 ns of the equilibrated system to determine any structural changes and differences in the number of interactions between the protein and drug at different time intervals.

### Principal component analysis

Principal component analysis (PCA) is a statistical technique that reduces the complexity of a data set in order to extract biologically relevant movements of protein domains from irrelevant localized motions of atoms. For PCA analysis the translational and rotational movements were removed from the system using gmx covar from GROMACS to construct a covariance matrix. Next the eigenvectors and eigenvalues were calculated by diagonalizing the matrix. The eigenvectors that correspond to the largest eigenvalues are called “principal components”, as they represent the largest-amplitude collective motions. We filtered the original trajectory and project out the part along the most important eigenvectors namely: vector 1 and 2 using gmx anaeig from GROMACS utilities. Furthermore, we visualized the sampled conformations in the subspace along the first two eigenvectors using gmx anaeig in a two-dimensional projection.

## Results

### Sequence and structural analysis

The amino acid sequence of HIV-1C IN shared 93.4% sequence identity with the template 5u1c amino acid sequence. The protein structure built for HIV-1C IN had a Global mean quality estimate score of 0.59 and 60% sequence similarity to 5u1c. The tetrameric structure of HIV-1C is shown in Fig 1, with each monomer labelled and colour coded. The VERIFY 3D score was 80.1%, the ERRAT overall quality score was 90% and higher for all four chains (A, B, C and D) and the Ramachandran plot indicated more than 90% of residues fell within the most favoured regions of the plot suggesting the predicted structure is a reliable model. Stability predictions indicated 15 RAMs to be destabilizing and five to be stabilizing for the protein structure based on SDM delta-delta G free energy scores (Table 1). The FoldX change in unfolded energy values indicated that the S140 was stabilizing, Q92 destabilizing and R143 neutral based on comparison with the WT structure each having values of 162.89, 131.94, 146.47 and 151.83 Kcal/Mol, respectively. Interaction analysis showed ten mutations resulted in a loss of polar contacts; three resulted in an increase in polar contacts, while seven showed no change in the number of polar contacts with neighbouring residues (Table 1).

**Fig 1.**
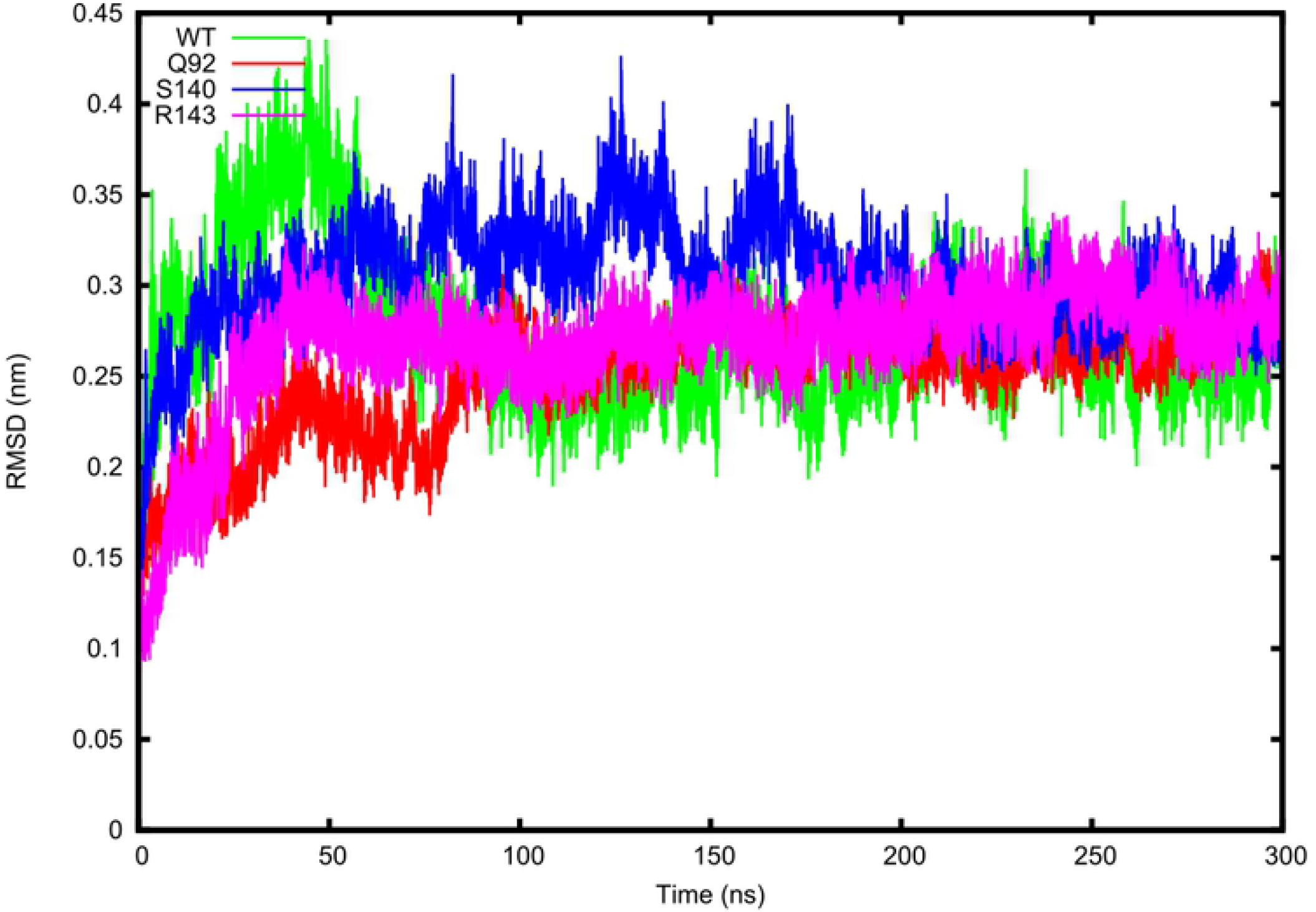
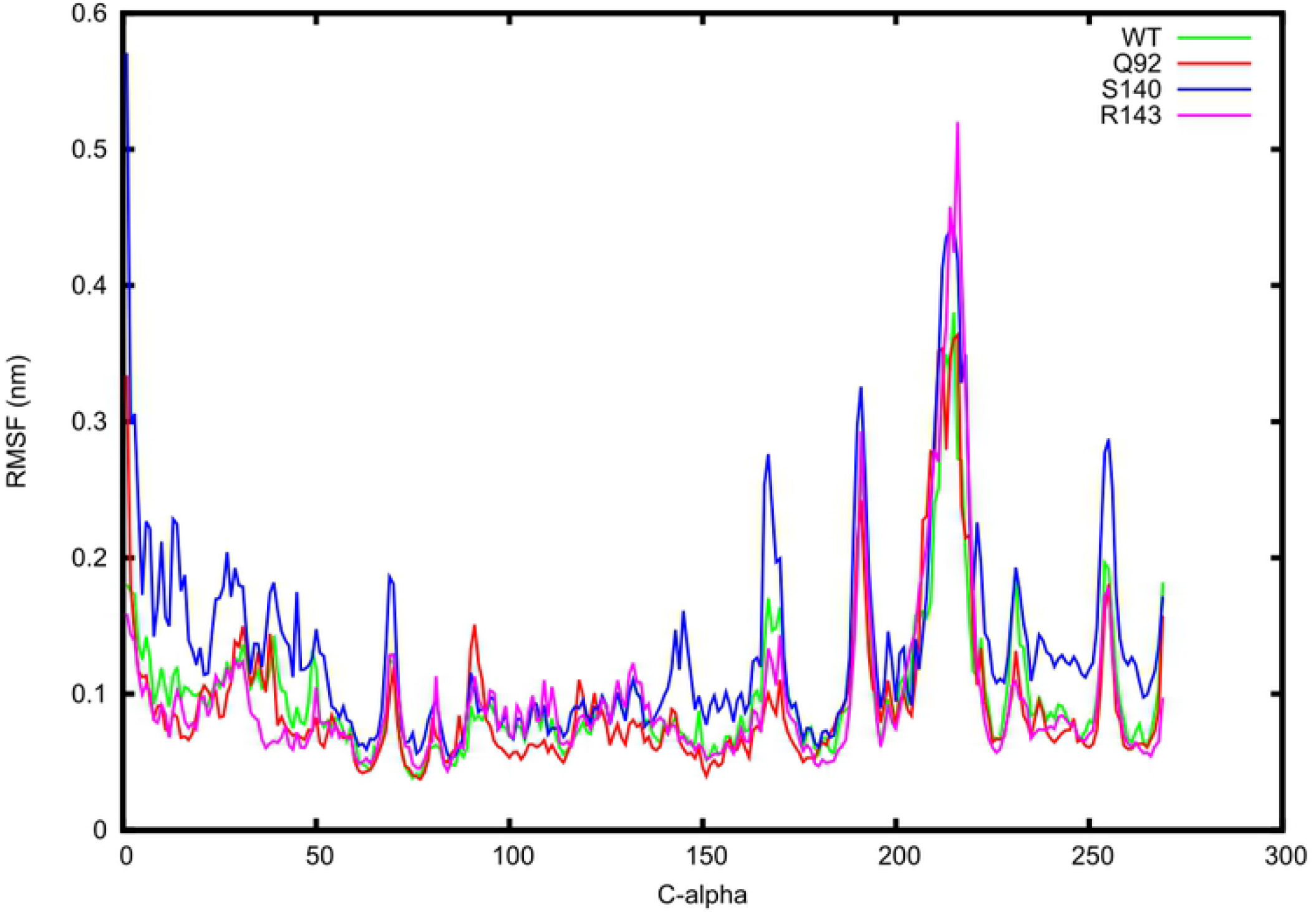
Tetrameric 3D structure of HIV-1C Integrase in complex with DNA, MG and drug Dolutegravir. Magnesium^2+^ ions (dirty violet spheres) are shown sitting in close proximity with Dolutegravir (brown) within the binding pocket (DDE motif) of the protein represented as navy blue sticks. A pair of dimers is seen encapsulating the viral/host DNA chimera, in which the two inner molecules (chain A and C) are shown in direct contact with DNA.

**Table 1.** Summary of stability predictions and polar interactions.

|  | SDM<sup>1</sup> | FoldX<sup>2</sup> |  | Polar Interactions |  |
| --- | --- | --- | --- | --- | --- |
| Mutation | Predicted ΔG (Kcal/Mol) | Total Energy ΔG (Kcal/Mol) | Energy Difference | Wild Type | Mutant |
| WT | N.A | 151.83 | N.A | N.A | N.A |
| T66A | -1.2 | 152.69 | 0.86 | 3 (H67, I73, ADE21) | 1 (I73) |
| T66I | 0.08 | 153.63 | 1.80 | 3 (H67, I73, ADE21) | 1 (I73) |
| T66K | -0.61 | 160.15 | 8.32 | 3 (H67, I73, ADE21) | 1 (I73) |
| E92Q | -0.16 | 131.94 | -19.89 | 3 (Q136, I113, T115) | 0 |
| E138K | -0.12 | 151.32 | -0.51 | 3 (Q136, I113, T115) | 2 (T115, I113) |
| E138A | -0.4 | 152.01 | 0.18 | 3 (Q136, I113, T115) | 2 (T115, I113) |
| E138T | 0.48 | 151.82 | -0.01 | 3 (Q136, I113, T115) | 2 (T115, I113) |
| G140S | -0.58 | 162.89 | 11.06 | 2 (T115, N117) | 3 (Q148, T115, N117) |
| G140A | -0.68 | 152.43 | 0.60 | 2 (T115, N117) | 2(T115, N117) |
| G140C | 0.39 | 154.81 | 2.98 | 2 (T115, N117) | 2(T115, N117) |
| Y143C | 0.14 | 152.49 | 0.66 | None | None |
| Y143R | -0.08 | 146.47 | -5.36 | None | 1 (S230) |
| Y143H | -0.07 | 152.28 | 0.45 | None | None |
| S147G | -0.18 | 151.16 | -0.67 | 2 (Q148, N144) | 1 (Q144) |
| Q148H | 0.63 | 157.78 | 5.95 | 3 (V151, P145, S147) | 2 (V151, P145) |
| Q148K | -0.78 | 151.33 | -0.50 | 3 (V151, P145, S147) | 3 (V151, P145, H114) |
| Q148R | -0.71 | 152.53 | 0.70 | 3 (V151, P145, S147) | 4 (D116,P145, V150, V151) |
| Q148N | -0.82 | 151.58 | -0.25 | 3 (V151, P145, S147) | 3 (V151, P145, H114) |
| N155H | -0.23 | 152.01 | 0.18 | 3 (V151, P145, S147) | 3 (E152, V151, K159) |
| R263K | -0.29 | 151.15 | -0.68 | 4 (Q146, N144, GUA18, ADE18) | 0 |

<sup>1</sup>negative values for $\Delta\Delta G$ indicate a stabilizing effect and positive values destabilizing.
<sup>2</sup>positive energy difference $\Delta G$ values $>1$ indicate a destabilizing effect, whereas values $1 \leq \Delta G \leq 0$ imply
a neutral effect and $\Delta G$ values $> -1$ indicate a stabilizing effect. Abbreviations used: N.A- not applicable.
The number in front of brackets is the total amount of interactions. Abbreviations of amino acids: A -
Alanine; D-Aspartic acid; E-Glutamic acid; G-Glycine; H-Histidine; I-Isoleucine; K-Lysine; N-
Asparagine; Q-Glutamine; R-Arginine; S-Serine; T-Threonine; Y-Tyrosine.

### Molecular dynamic simulations

All the MD trajectory analysis considered the single chain A (monomer) of the IN protein in contact with the drug DTG. Trajectory analysis of the RMSD of the backbone indicated that the WT system reached equilibrium after 100 ns as well as the Q92, R143 and S140 mutant systems (Fig 2A). Only S140 showed higher RMSD fluctuation values compared to the WT, R143 and Q92 systems (Fig 2A). RMSF analysis clearly showed higher flexibility for the S140 mutant system, with four highly flexible regions (residues 68 - 70, 142 - 146, 166 - 170 and 253 - 256) compared to the WT, Q92 and R143 systems (Fig 2B). These flexible regions affect the 140’s loop region that regulates drug binding. The Radius of gyration values indicated decreasing values for R143 and E92 compared to the WT and S140 mutant system (Fig 2C). Plotting the first two principal components provided insight into the collective movement of each protein atom. The 2D projections of the first and second principal components for the WT, Q92, R143 and S140 systems are shown in Figure 2D. Calculation of the covariance matrix values after diagonalization showed a significant increase for the S140 system (18.33 nm) compared to the other three systems WT, Q92 and R143 each having 9.58 nm, 8.98 nm and 10.41 nm lower values, respectively. Distance analysis indicated a smaller average distance of 0.21 ± 0.01 nm and 0.22 ± 0.01 nm between the WT, R143 MG ion and drug DTG systems compared to Q92 and S140 each having a distance of 0.41 ± 0.04 nm, 0.99 ± 0.20 nm, respectively (Fig 2D).

**Fig 2.**
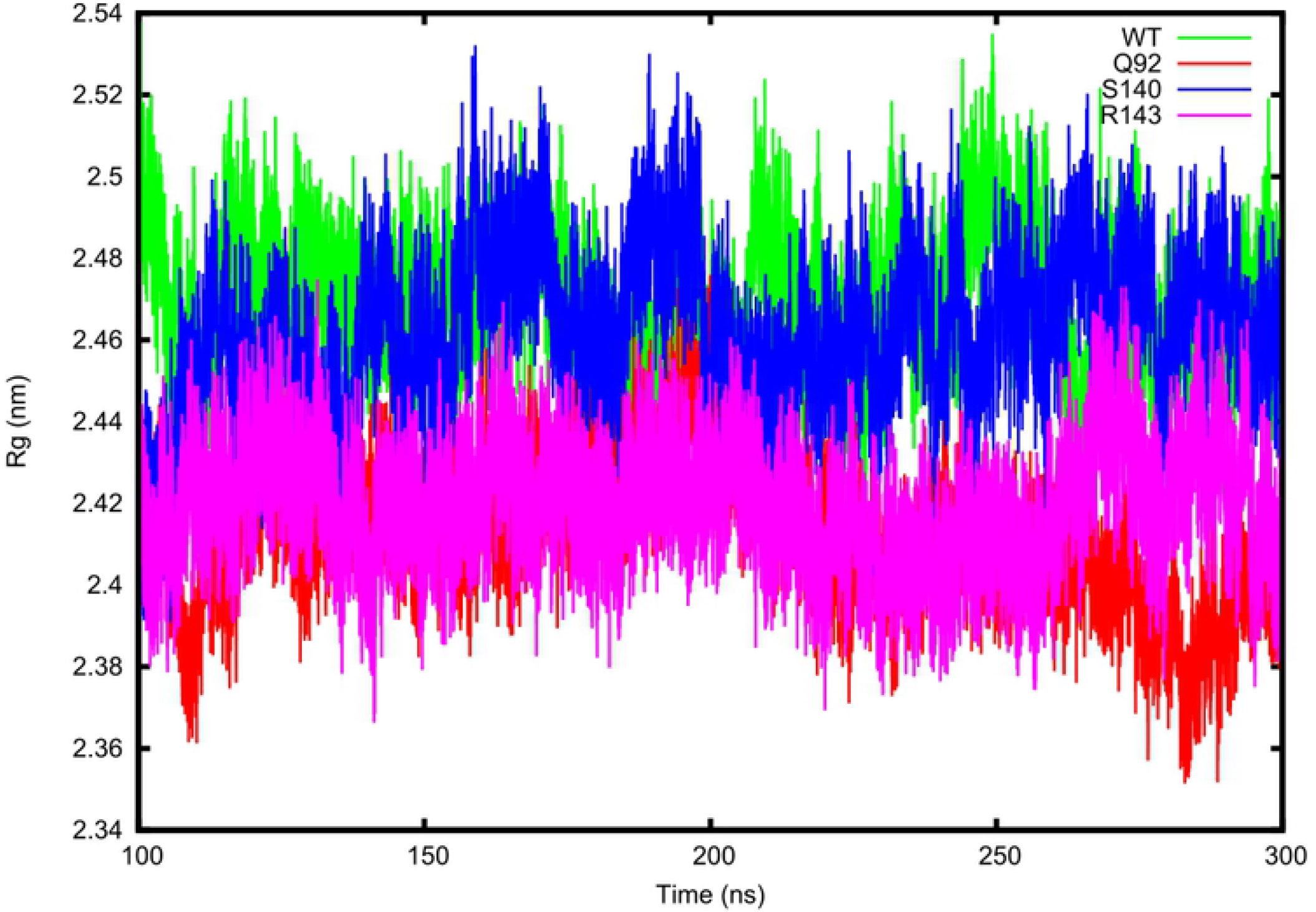
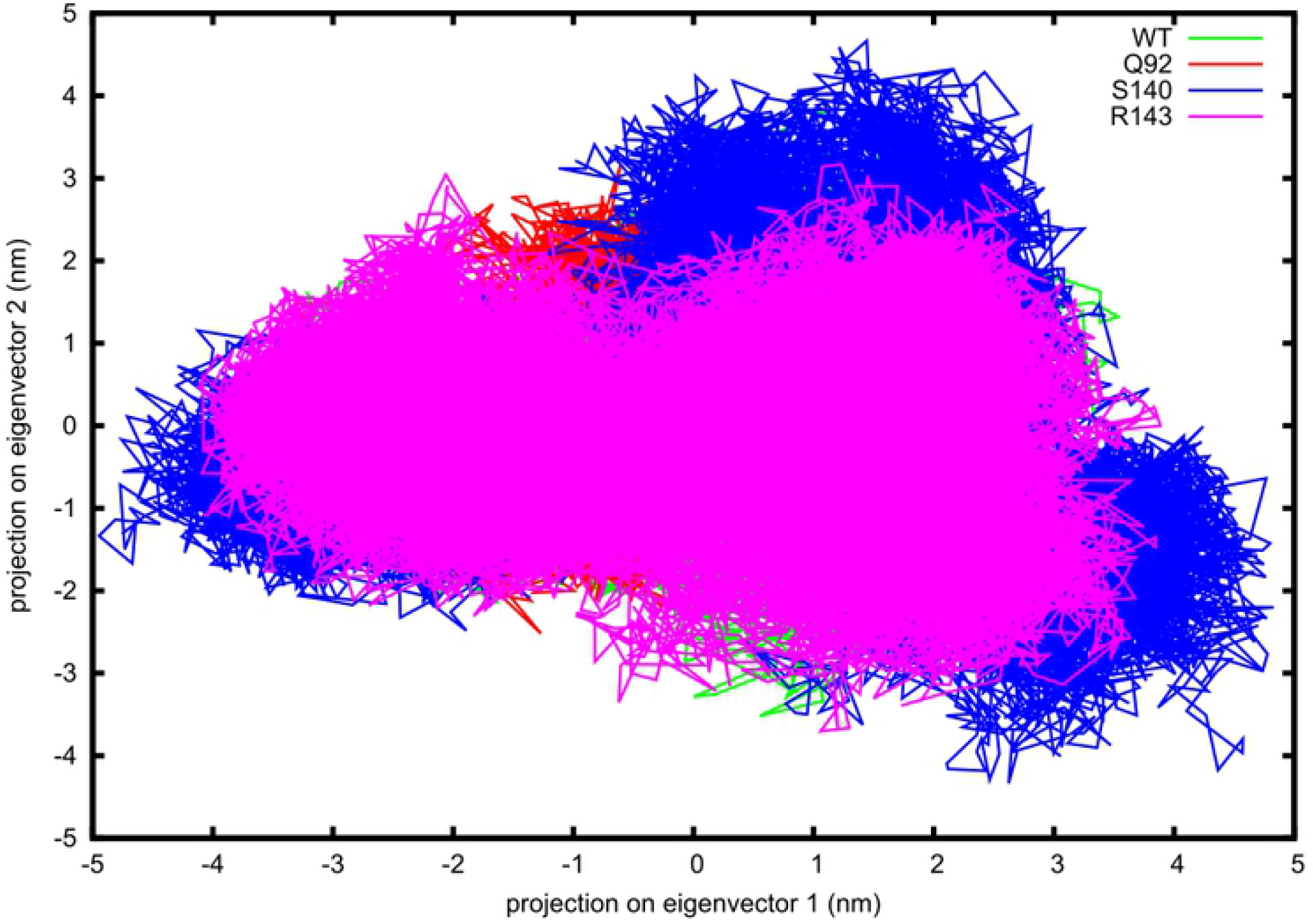
Trajectory analysis of the four simulation systems. (A) Change in backbone RMSD for the WT, Q92, S140 and R143 systems plotted over 300 ns. (B) Change in RMSF for the C-alpha residues for the WT, Q92, S140 and R143 systems plotted over the last 200ns. (C) Measure of compactness for the WT, Q92, S140 and R143 systems plotted over the last 200 ns. (D) First two principal components plotted for the WT, Q92, S140 and R143 systems plotted over the last 200 ns.

### Interaction analysis

We performed interaction analysis for five snapshots (every 50 ns) of each of the simulation systems to determine which residues played a role in the binding of DTG to the protein in the WT and mutant protein structures. For the WT system, interactions were observed between known active site residues D64, D116 and N148, MG ion and also to DNA nucleotides (Table 2). Similarly, interactions were observed between known active site residues D64, R143, N148, MG ion and DNA nucleotides for the R143 system (Table 2). Interestingly, no active site residue interactions between DTG and S140 structure was observed and no MG ionic interactions for S140. Only the Q92 system showed interactions with active site residues but no MG ionic interactions (Table 2). Figs A-D in S1 Supporting Information File shows the relative location of the drug to the MG ion and active site for snapshot 1 taken at 100 ns. Compared to the WT, R143, Q92 the drug DTG is located close to the MG ion and active site while for S140 the drug is located outside the catalytic site suggesting the mutation is able to expel the drug from the binding pocket over time (Figs A-D in Supporting Information File).

**Table 2.** Summary of interaction analysis.

| Structure | Cluster | Interactions |  |
| --- | --- | --- | --- |
|  |  | Hydrogen bonds | Ionic |
| WT | 1 (100 ns) | 2 (GUA22<sup>a</sup>, D116) | MG |
|  | 2 (150 ns) | 3 (THY11<sup>a</sup>, D64, D116) | MG |
|  | 3 (200 ns) | 2 (GUA22<sup>a</sup>, D116) | MG |
|  | 4 (250 ns) | 4 (THY11<sup>a</sup>, GUA22<sup>a</sup>, D64, D116) | MG |
|  | 5 (300 ns) | 2 (THY11<sup>a</sup>, N148) | MG |
| R143 | 1 (100 ns) | 4 (THY11<sup>a</sup>, GUA22<sup>a</sup>, D64, N148) | MG |
|  | 2 (150 ns) | 4 (GUA22<sup>a</sup>, D64, R143, N148) | MG |
|  | 3 (200 ns) | 4 (GUA22<sup>a</sup>, D64, R143, N148) | MG |
|  | 4 (250 ns) | 4 (GUA22<sup>a</sup>, D64, R143, N148) | MG |
|  | 5 (300 ns) | 5 (THY11<sup>a</sup>, GUA22<sup>a</sup>, GUA22<sup>a</sup>, D64, N148) | MG |
| Q92 | 1 (100 ns) | 3 (CYT20<sup>a</sup>, D116, P145) | None |
|  | 2 (150 ns) | 3 (D116, P145, E152) | None |
|  | 3 (200 ns) | 3 (CYT20<sup>a</sup>, H21, D116) | None |
|  | 4 (250 ns) | 4 (CYT20<sup>a</sup>, P142, P145, E152) | None |
|  | 5 (300 ns) | 3 (CYT20<sup>a</sup>, P145, N148) | None |
| S140 | 1 (100 ns) | 3 (GUA22<sup>a</sup>, ADE25<sup>a</sup>, ADE27<sup>a</sup>) | None |
|  | 2 (150 ns) | 3 (GUA22<sup>a</sup>, ADE25<sup>a</sup>, ADE27<sup>a</sup>) | None |
|  | 3 (200 ns) | 3 (GUA22<sup>a</sup>, ADE25<sup>a</sup>, ADE27<sup>a</sup>) | None |
|  | 4 (250 ns) | 3 (THY11<sup>a</sup>, GUA22<sup>a</sup>, ADE27<sup>a</sup>) | None |
|  | 5 (300 ns) | 3 (THY11<sup>a</sup>, GUA22<sup>a</sup>, ADE27<sup>a</sup>) | None |

<sup>a</sup>Interactions with DNA nucleotides.
Abbreviations of DNA nucleotides: ADE-Adenine; CYT-Cytosine; GUA-Guanine; THY-Thymine. Abbreviations of amino acids: D-Aspartic Acid; E-Glutamic Acid; H-Histidine N-Asparagine; P-Proline; R-Arginine.

## Discussion

Sequence variation between HIV-1B and HIV-1C can have a meaningful contribution to the protein structure, affecting protein drug interaction. We investigated known drug RAMs associated with RAL, EVG and DTG resistance to determine their effect on the protein structure and drug binding. The structural modelling of HIV-1C IN considered a homologous template of high sequence identity, and good overall target sequence coverage, compared to previous homology models that considered templates of low sequence identity. We could therefore accurately reconstruct HIV-1C using the close homolog HIV-1B crystal structure as template to infer accurate drug interactions. Further inspection of the overall structure confirmed accurate prediction of more than 90% of domains within the protein structure, compared to the template HIV-1B structure. The quality analysis provided support for the predicted model based on side chain conformations. Stability predictions showed contrasting results to interaction analysis, whereby amino acid substitutions that resulted in a gain of interactions was predicted to be destabilising. The FoldX changes in energy values were similar to interaction analysis for the three mutant structures under investigation. To fully comprehend the effects of individual mutations we opted to use molecular dynamic (MD) simulations to understand the effect of selected mutations on protein movement and drug interactions. MD analyses have shown to be successful in quantifying small changes in protein structures that can affect overall drug binding [23]. We randomly selected three mutations Q92, S140 and R143 to investigate the role of amino acid substitutions on protein structure and drug binding in comparison to the WT protein structure. Analysis of the change in trajectory of the mutant systems compared to the wild type suggested less stability and higher fluctuation of the G140S mutant system compared to the WT system. We also confirmed the destabilizing effect of the G140S mutant using principal component analysis which suggested larger randomized concerted movement for the G140S mutant compared to the WT, Q92 and R143 systems. These findings are similar to Chen et al. [10] which showed that the G140S mutations can either stabilize or destabilize the 140’s loop region. In our case, the 140’s loop region is stabilized by the G140S mutation that could prevent drug binding. This is supported by pairwise distance analysis confirming a larger distance between the MG ion and drug DTG for the G140S mutant system compared to the WT and R143. Further interaction analysis was performed to confirm the hypothesis that the S140 mutation could prevent drug binding by extracting structures at different snapshots of the simulation. Here, we found that the G140S mutation resulted in the drug being expelled from the binding pocket. We also observed weaker interactions for the Q92 mutation but stronger interactions for R143 mutant. The findings from our study suggest that patients should first be screened for mutations and or novel variants to determine if the drug DTG will be efficacious or not. The model generated in this study can be used to tease out the effects of novel variants. A few limitations of this study are the use of RAL, EVG mutants and not considering novel RAL or DTG mutations. We also only simulated single mutations instead of double however we have yet to identify double mutants within the South African cohort of HIV-1C infected patients. Future work will include viral fitness assays to determine the effect of the mutant S140 on the HIV-1C virus and the effect of the drug DTG on the virus ability to persist in the presence of the drug.

## Acknowledgements

We would like to acknowledge the Poliomyelitis Research Foundation (PRF) of South Africa for funding this project. The Centre for High Performance Computing (CHPC), Rondebosch, South Africa for access to their high performance computing cluster and the Higher Education Department, next Generation of Academic Programme (nGAP) for full time academic positions of Dr Cloete and Dr Jacobs. This study was funded by The South African Research Chairs Initiative of the Department of Science and Technology (DST), the National Research Foundation (NRF) of South Africa, and the South African Medical Research Council (SAMRC).

## Supporting information

**S1 Fig.**
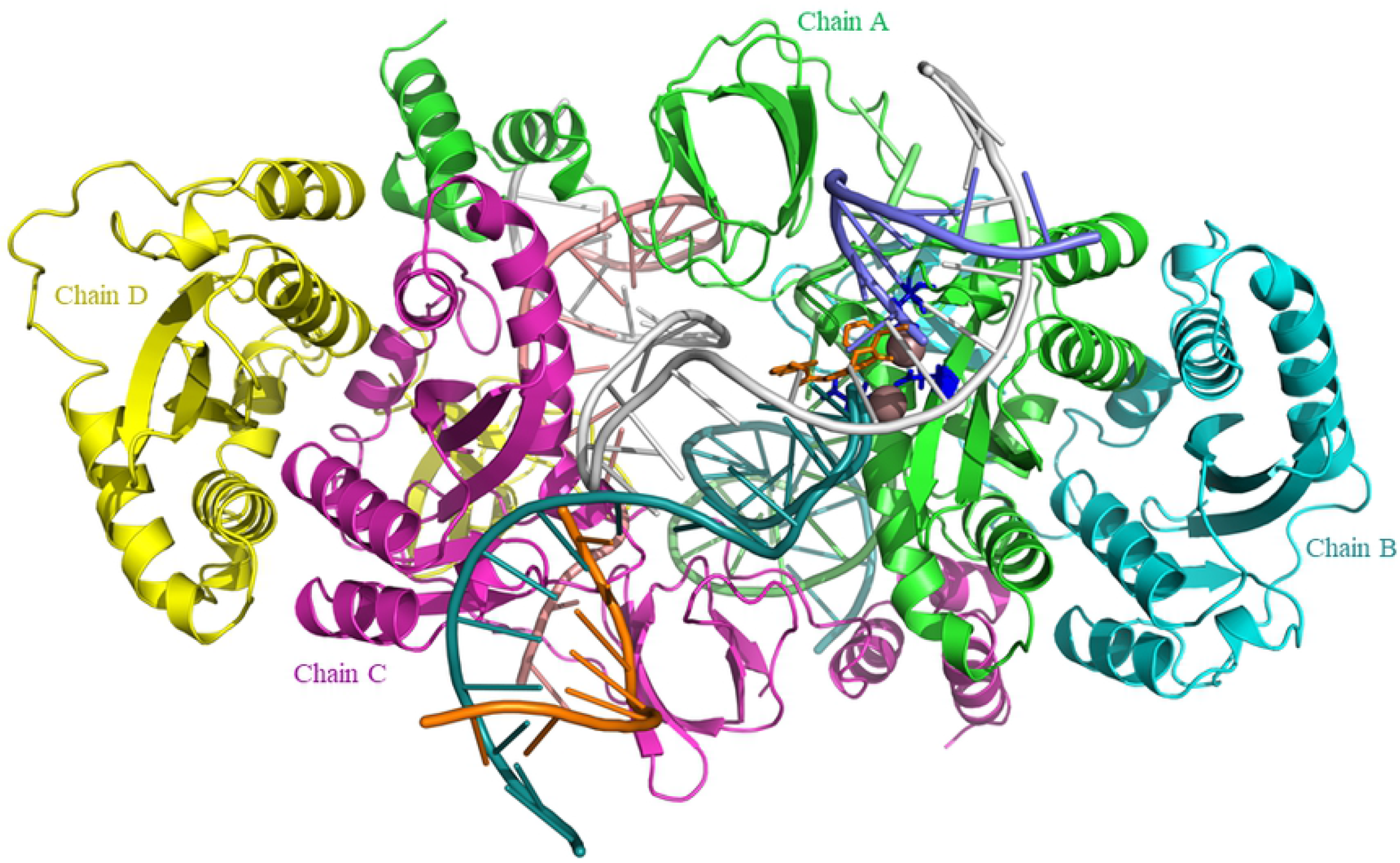
Clustering analysis for the four simulation systems. (A) Cartoon representation of the WT HIV-1C integrase structure showing the location of the drug taken at 100 ns. (B) Cartoon representation of the R143 HIV-1C integrase structure showing the location of the drug taken at 100 ns. (C) Cartoon representation of the Q92 HIV-1C integrase structure showing the location of the drug taken at 100 ns. (D) Cartoon representation of the S140 HIV-1C integrase structure showing the location of the drug taken at 100 ns.

